# Ranked Choice Voting for Representative Transcripts with TRaCE

**DOI:** 10.1101/2020.12.15.422742

**Authors:** Andrew J. Olson, Doreen Ware

## Abstract

**Summary:** Genome sequencing projects annotate protein-coding gene models with multiple transcripts, aiming to represent all of the available transcript evidence. However, downstream analyses often operate on only one representative transcript per gene locus, sometimes known as the canonical transcript. To choose canonical transcripts, TRaCE (Transcript Ranking and Canonical Election) holds an ‘election’ in which a set of RNA-seq samples rank transcripts by annotation edit distance. These sample-specific votes are tallied along with other criteria such as protein length and InterPro domain coverage. The winner is selected as the canonical transcript, but the election proceeds through multiple rounds of voting to order all the transcripts by relevance. Based on the set of expression data provided, TRaCE can identify the most common isoforms from a broad expression atlas or prioritize alternative transcripts expressed in specific contexts.

**Availability and Implementation:** Transcript ranking code can be found on GitHub at {{https://github.com/warelab/TRaCE}}

**Supplementary information:** Additional data are available in the github repository.

## Introduction

Genome sequencing projects often use complex, automated annotation pipelines to build reference sets of gene models. These pipelines mask repeats in the assembled genome, align protein and transcript evidence, and build gene models by aggregating overlapping alignments that adhere to known or inferred splice site patterns (Hoff et al. 2019; Campbell et al. 2014; Haas et al. 2003). Before a project releases a set of high-confidence gene models, additional filtering steps may remove transcript models that lack homology or are subject to nonsense-mediated degradation (NMD).

Alternative splicing contributes to the functional diversity of a genome (Black 2003); and new sequencing technology such as PacBio IsoSeq can capture splice variants at an unprecedented scale (Wang et al. 2016; Zhang et al. 2019; Bruijnesteijn et al. 2018). However, this heightened sensitivity can lead to the detection of non-functional transcripts, which can be misreported by gene builders as biologically relevant splice variants. Furthermore, it is possible for partially processed transcripts containing retained introns that neither disrupt the reading frame nor introduce stop codons to be promoted to canonical transcripts (Figure 1).

**Figure 1.**
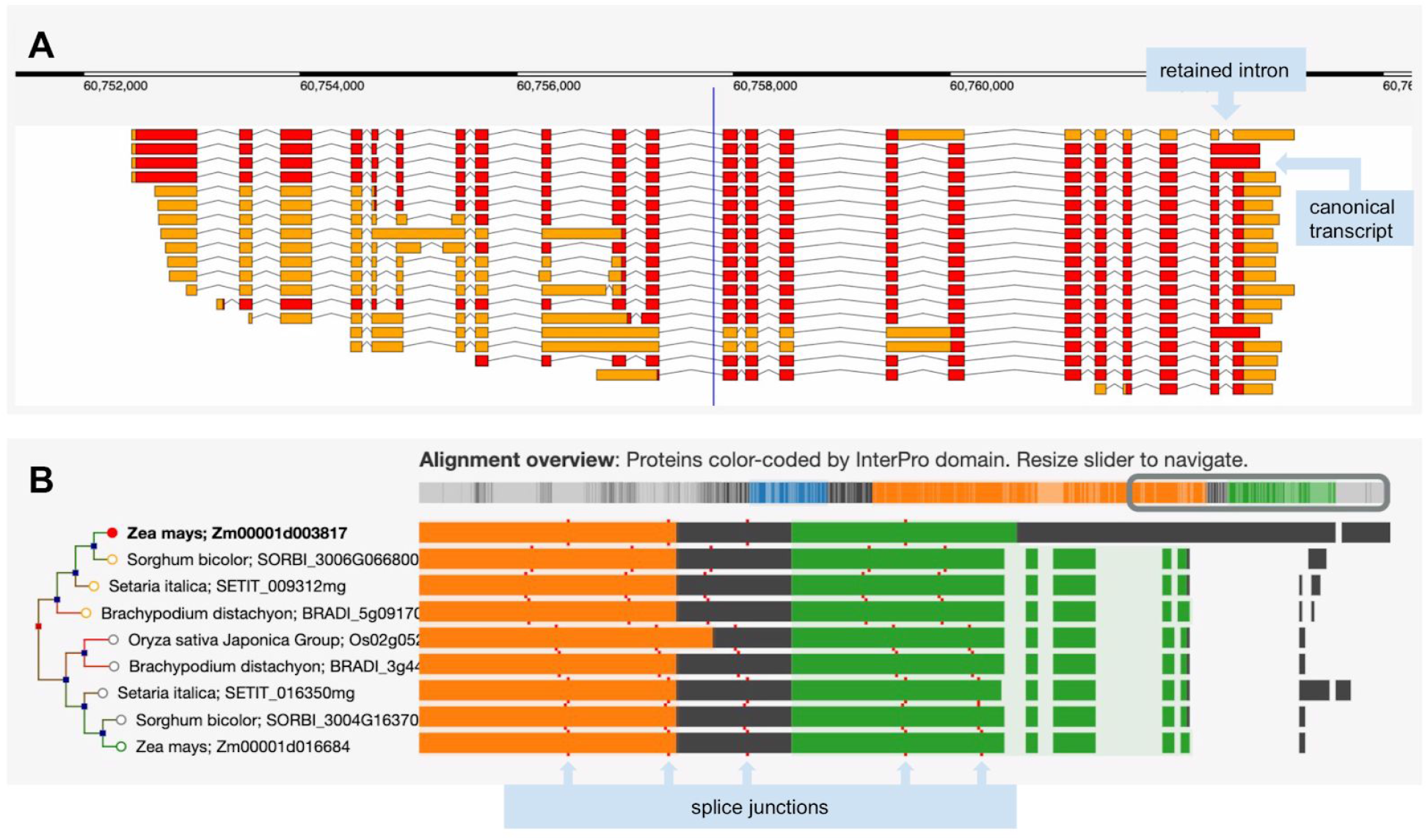
**A)** The complex set of transcript models for the *Zea mays* B73 gene sbe4 (starch branching enzyme4). Red blocks show the predicted coding regions, and orange blocks are untranslated regions. The longest translation contains a retained intron and was selected as the canonical transcript for Compara gene tree analysis. **B)** The left side shows a portion of the gene tree focused on this maize gene and displaying homologs from *Sorghum bicolor, Setaria italica, Brachypodium distachyon,* and *Oryza sativa Japonica*. The right side shows regions of protein sequences participating in the multiple sequence alignment, color coded by InterPro domain. The first row shows a unique region relative to other species that derives from the retained intron.

Comparative gene tree analysis platforms such as Ensembl Compara (Herrero et al. 2016) operate on a single canonical transcript for each gene locus. In the absence of a curated canonical transcript, this is usually defined as the longest transcript with the longest translation, but this definition does not necessarily select the best representative transcript for a gene locus. Subsequently developed techniques have defined canonical isoforms based on expression level, sequence conservation, annotation of functional domains, or some combination of these features (Li et al. 2014; Rodriguez et al. 2018; The UniProt Consortium et al. 2016).

We developed TRaCE (Transcript Ranking and Canonical Election) to choose canonical transcripts based on data typically available at the time of a new genome annotation. In this approach, transcripts are ranked according to length, domain coverage, and according to how well they represent a diverse population of transcriptome RNA-seq data. An ‘election’ based on ranked-choice voting selects a canonical transcript that is the first- or second-choice transcript for the majority of samples. The election proceeds through multiple rounds, effectively sorting all transcripts by relevance. Here we present the TRaCE algorithm and results obtained by running TRaCE on *Zea mays* and *Homo sapiens* gene annotations. In addition, we describe validation of TRaCE predictions by manual curation (Tello-Ruiz et al. 2019) and comparison to APPRIS (Rodriguez et al. 2018) human transcript classifications.

## Methods

The first step in preparing to run TRaCE is to gather a diverse set of RNA-seq expression data covering a wide variety of tissues or conditions to act as ‘voters’ in the upcoming elections. The next step is to align the reads, assemble sample-specific transcripts, and quantify their expression. Each reference gene model with multiple transcripts (candidates) will hold an election to sort the reference transcripts by relevance (Figure 2).

**Figure 2.**
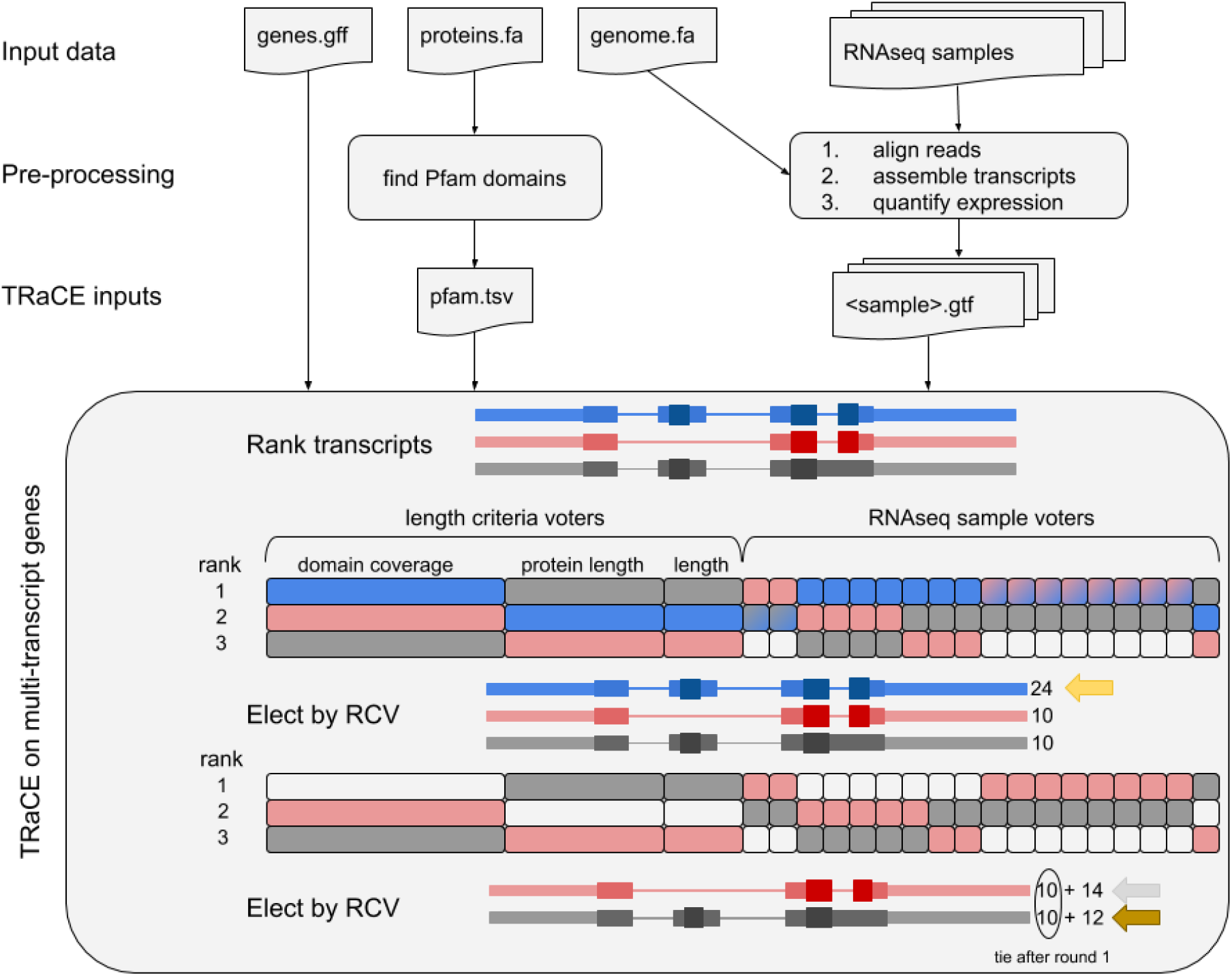
Flowchart of preparation of TRaCE inputs and a schematic of the rank-choice voting (RCV) approach to select transcripts for an example gene with three transcripts (blue, red, gray). Exon thickness corresponds to non-coding, coding, and functional regions with Pfam domains. Voters are represented by rectangles, and rank transcripts by length criteria (9, 6, or 3 votes) or AED (1 vote per sample). Eight of the samples rank the red and blue transcripts equally (blue-red gradient), so both get tallied in round 1. RCV selects the blue transcript first with 24 rank 1 votes. After removing the blue votes from consideration, the red and gray transcripts tie with 10 rank 1 votes, but the red transcript is elected with 14 rank 2 votes.

In each election, samples rank the candidate transcripts using a modified annotation edit distance (AED) comparing them to the most highly expressed overlapping sample-specific transcripts (Eilbeck et al. 2009). AED ranges from 0 (perfect agreement) to 1 (no overlap). Because there may be insufficient data to assemble full-length transcripts from samples in which the gene is expressed at low levels, the AED score is restricted to overlapping portions of candidate transcripts. A maximum AED score cutoff (default, 0.5) prevents samples from voting for candidate transcripts with very little similarity. There are also cutoff parameters for minimum expression level (default TPM, 0.5) and proportion overlapping (default, 0.5) to filter out some noise in the sample transcriptome data. The election includes additional voters that rank transcripts based on domain coverage, protein length, and transcript length. To avoid overwhelming the length-based voters when running TRaCE with many samples, sample votes are weighted to balance the electorate.

Once each sample and the length based voters have ranked the transcripts, the election proceeds in multiple rounds selecting winners until no candidates remain. In each round, TRaCE tallies votes for top-ranked candidates; and so long as there is a tie for first place, votes for the subsequent rankings are added to the tally.

## Results

We ran TRaCE on a pre-release set of *Zea mays* B73 gene models with the set of 10 RNA-seq samples that had already been aligned to the genome as part of the evidence-based gene annotation pipeline (Hufford et. al., in preparation). The samples were derived from shoot, root, embryo, endosperm, ear, tassel, anther, and three leaf sections (base, middle, and tip). StringTie version 1.3.5 (with the --rf flag) was used for transcript assembly and quantification (Pertea et al. 2016) and InterProScan version 5.38-76.0 was run to identify Pfam domains (Mulder and Apweiler 2007). The *Zea mays* B73 V5 annotation set (Zm00001eb) has 15,162 multi-transcript protein-coding gene models; for 5,616 of these (37%), the canonical transcript chosen by TRaCE was not the longest isoform. TRaCE selected canonical transcripts for the genome annotations of 25 additional maize accessions, 33-38% of which were not the longest isoform (Suppl Table 1).

We also ran TRaCE on human GRCh38 annotations (Frankish et al. 2019) with a diverse panel of 127 samples of human RNA-seq data covering the development of seven major organs (brain, cerebellum, heart, kidney, liver, ovary and testis) from 4 weeks post-conception to adulthood (https://www.ebi.ac.uk/gxa/experiments/E-MTAB-6814/Results). Reads were aligned with hisat2 version 2.1.0 (--dta --reorder), transcripts were assembled and quantified with stringtie version 2.1.4 (--conservative), and protein-coding reference transcripts were annotated with Pfam domains using InterProScan version 5.38-76.0 (Pertea et al. 2016; Mulder and Apweiler 2007). The GRCh38 annotation set has 15,984 multi-transcript protein-coding gene models, 4,458 (28%) of which had a TRaCE canonical that was not the longest isoform. For comparison, the principal isoform according to APPRIS was not the longest isoform in 3,824 (24%) of the multi-transcript gene models.

## Validation

We used two approaches to validate TRaCE’s predictions. First, we modified an interactive gene tree viewer, designed to flag problematic gene models by visual inspection of the multiple sequence alignment and domain annotations (Tello-Ruiz et al. 2020). We used this interface to compare maize B73 V5 canonical transcripts (Zm00001eb) selected by TRaCE with the prior set of maize V4 canonical transcripts (Zm00001d) selected by length criteria alone. A random selection of 173 pairs of genes for which the TRaCE canonical was not the longest transcript were evaluated in the gene tree viewer and flagged if the alignment was inconsistent with outgroup orthologs. Genes were flagged if there was a relative gain or loss of conserved sequence within the transcript or at either end. Of these gene pairs, 32% were flagged as problematic in Zm00001d only, 4% in Zm00001eb only, and 5% in both versions (Suppl Table 2). The most common issue in the flagged Zm00001d gene models was gain of sequence due to an intron retention. Thus, according to this approach, TRaCE was selecting better-conserved isoforms than the prior length-based algorithm.

In the second approach, TRaCE predictions were also validated by student curators who were given a subset of 48 gene models with two to five predicted transcripts each for which TRaCE’s top-ranked isoform was not the longest isoform. The students, who were not aware of TRaCE’s output, were asked to rate transcripts as best, good, or poor, based on viewing the gene structure and expression evidence in the Apollo genome browser (Dunn et al. 2019). Each gene model was curated by at least 3 different students. The transcript ratings were mapped to a score (best 2, good 1, poor −1). Transcript rankings from TRaCE and rankings based on length alone were compared to rankings based on curator scores. For each rank (1-5), we calculated the sum of the curator scores for the associated transcripts. The correlation between the length based ranking and the curator based ranking was 0.917, whereas the TRaCE and curator rankings had a higher correlation coefficient of 0.985 (Suppl Table 3).

To test the performance of TRaCE on human transcript annotations, we selected a comparable subset of 2,029 genes with two to five protein-coding transcripts each, for which the TRaCE winner was not the longest transcript. We used a similar scoring method to compare TRaCE rankings to rankings by length alone based on APPRIS transcript classifications (Principal 2, Alternative 1, Minor −1). The length-based and APPRIS-based scores had a correlation coefficient of 0.788, whereas the TRaCE and APPRIS rankings had a correlation coefficient of 0.966 (Suppl Table 3). Given that TRaCE and APPRIS perform very similarly when selecting canonical transcripts, we are confident that TRaCE represents a suitable alternative that could be applied easily to new and existing genome annotations.

## Acknowledgements

The authors thank Ware lab summer students Sarina Awatramani and Pragati Muthukumar for their preliminary research on intron retentions in Zm00001d, collaborators on the maize NAM project for feedback on TRaCE selected canonical transcripts, Kevin Ahern and students from his genetics course at Cornell University for curation, Bruno Contreras-Moreira for suggesting a comparison with APPRIS, Cristina Marco and Marcela K. Tello-Ruiz for running curation jamborees and testing the algorithm, Liya Wang, Marcela K. Tello-Ruiz, and Christopher Patil for reviewing and editing the manuscript and Augusto Diniz for the TRaCE acronym.

